# Development of genetic tools for the thermophilic filamentous fungus *Thermoascus aurantiacus*

**DOI:** 10.1101/2020.06.09.143198

**Authors:** Raphael Gabriel, Julia Prinz, Marina Jecmenica, Carlos Romero-Vazquez, Pallas Chou, Simon Harth, Lena Floerl, Laure Curran, Anne Oostlander, Linda Matz, Susanne Fritsche, Jennifer Gorman, Timo Schuerg, André Fleißner, Steven W. Singer

**Affiliations:** Biological Systems and Engineering Division, Lawrence Berkeley National Laboratory, 5885 Hollis Street, Emeryville, CA 94608, USA; Joint BioEnergy Institute, Emeryville, CA 94608, CA, USA; Institut für Genetik, Technische Universität Braunschweig, Spielmannstr. 7, 38106 Braunschweig, Germany; Department of Applied Genetics and Cell Biology, University of Natural Resources and Life Sciences Vienna (BOKU), Muthgasse 18, 1190 Vienna, Austria; Austrian Centre of Industrial Biotechnology (ACIB), Muthgasse 11, 1190 Vienna, Austria; Department of Biotechnology, University of Natural Resources and Life Sciences (BOKU), Muthgasse 18, 1190 Vienna, Austria; University of Puerto-Rico, Rio Piedras, San Juan, Puerto Rico 00925; American High School, 36300 Fremont Blvd, Fremont, CA 94536; Frankfurt Institute of Molecular Biosciences, Goethe University Frankfurt, 60438, Frankfurt am Main, Germany; École Polytechnique Fédérale de Lausanne, Lausanne, Vaud, 1015, Switzerland; Department of Food Science and Technology, University of Natural Resources and Life Sciences Vienna (BOKU), Muthgasse 18, 1190 Vienna, Austria

**Keywords:** Filamentous fungi, *Thermoascus aurantiacus*, *Agrobacterium tumefaciens*, genetic transformation, CRISPR/Cas9 system, sexual crossing, xylanases, enzyme production

## Abstract

**Background:** Fungal enzymes are vital for industrial biotechnology, including the conversion of plant biomass to biofuels and bio-based chemicals. In recent years, there is increasing interest in using enzymes from thermophilic fungi, which often have higher reaction rates and thermal tolerance compared to currently used fungal enzymes. The thermophilic filamentous fungus *Thermoascus aurantiacus* produces large amounts of highly thermostable plant cell wall degrading enzymes. However, no genetic tools have yet been developed for this fungus, which prevents strain engineering efforts. The goal of this study was to develop strain engineering tools such as a transformation system, a CRISPR/Cas9 gene editing system and a sexual crossing protocol to improve enzyme production.

**Results:** Here we report *Agrobacterium tumefaciens-*mediated transformation (ATMT) of *T. aurantiacus* using the *hph* marker gene, conferring resistance to hygromycin B. The newly developed transformation protocol was optimized and used to integrate an expression cassette of the transcriptional xylanase regulator *xlnR*, which led to up to 500% increased xylanase activity. Furthermore, a CRISPR/Cas9 gene editing system was established in this fungus, and two different gRNAs were tested to delete the *pyrG* orthologue with 10% and 35% deletion efficiency, respectively. Lastly, a sexual crossing protocol was established using a hygromycin B- and a 5-fluororotic acid-resistant parent strain. Crossing and isolation of progeny on selective media was completed in a week.

**Conclusion:** The genetic tools developed for *T. aurantiacus* can now be used individually or in combination to further improve thermostable enzyme production by this fungus.

## Background

Due to the potentially deleterious impacts of climate change, which is mainly caused by the use of fossil resources, great efforts have been made to explore the applicability of lignocellulosic plant biomass as sustainable alternative to fossil fuels. Lignocellulosic biomass is the most abundant organic material on earth, consisting primarily of the sugar polymers cellulose and hemicellulose and the aromatic polymer lignin [1,2]. These sugar polymers can be deconstructed by enzymes (cellulases and hemicellulases) into simple sugars that can be further converted into biofuels and other bio-based products using metabolically engineered bacterial and fungal hosts, which will reduce our dependence on finite fossil resources [3]. The cost-efficient deconstruction of lignocellulose is currently the largest obstacle preventing biofuels from becoming competitive with fossil fuels.

Filamentous fungi are very effective lignocellulose degraders, possessing an arsenal of secreted enzymes that digest cellulose and hemicellulose [4]. These organisms have evolved an elaborate sensing system to detect the components of lignocellulosic biomass and fine-tune expression of cellulase and hemicellulase genes [5]. Therefore, filamentous fungi are the most important industrial cellulase producers [6,7].

Recently, there is increased interest in establishing thermophilic organisms that secrete thermostable enzymes for the conversion of plant biomass to biofuels [8–10]. The thermophilic fungi *Thielavia terrestris* and *Myceliophthora thermophila* produced enzymes that were more active across all temperatures tested and released more sugars from pretreated plant biomass than the enzymes of the mesophiles *Trichoderma reesei* and *Chaetomium globosum* [11]. In a separate study, enzymes from another thermophilic fungus, *Thermoascus aurantiacus*, demonstrated a higher level of sugar release from ionic liquid pretreated switchgrass than *T. terrestris* enzymes and showed activities comparable to the commercial enzymatic mixture CTec2 [12].

*T. aurantiacus* is a homothallic fungus that grows optimally at 50°C. Induction experiments indicated that both cellulases and xylanases were induced by controlled feeding with xylose, suggesting that the regulatory systems for enzyme expression in *T. aurantiacus* had similarities to the regulatory system in *Aspergillus niger* [13]. These initial results make *T. aurantiacus* an intriguing host for thermostable enzyme production. Improving enzyme production and investigating regulation of cellulase and xylanase expression in *T. aurantiacus* is limited by the absence of genetic tools for this promising fungus.

Efficient strain engineering requires genetically tractable hosts. Several methods have been established to genetically engineer filamentous fungi such as: protoplast transformation, electroporation, biolistics and *Agrobacterium tumefaciens*-mediated transformation (ATMT) [14]. ATMT relies on the ability of the plant pathogen *A. tumefaciens* to inject DNA into plant cells and other eukaryotic cells. In this manner, various genetic modifications have been made in fungal genomes, including applying CRISPR/Cas9-based gene editing systems [15]. The initial development of the CRISPR/Cas9 system for filamentous fungi often involved the deletion of counter-selectable marker genes such as *pyrG* [16] and *amdS* [17], which allows the fungus to grow in the presence of otherwise toxic 5-fluororotic acid or fluoroacetamide, respectively. Sexual crossing is another versatile tool, which accelerates strain engineering through combining desired phenotypes, mapping genomic loci, removing undesired mutations and generating genetically uniform fungal homokaryons [18,19]. Notably, sexual crossing is not possible with a variety of industrially highly relevant fungi, and a sexual cycle was only recently established for the classic cellulase producer *T. reesei* [20–22].

Genetic tools have been successfully applied to generate high enzyme secreting strains. A *Penicillium oxalicum* strain with strongly increased cellulase production was generated through overexpression of *clrB* and deletion of the cellulase repressors *creA* and *bglR* [23]. This strain displayed equivalent enzyme production compared to the industrial cellulase hypersecreting *P. oxalicum* strain JU-A10-T, which was generated through classical mutagenesis. Increases in cellulase and xylanase secretion were also achieved through overexpression of x*lnR* and *clrB* and deletion of *creA* in this fungus [24]. Similarly, a *M. thermophila* cellulase hypersecreting strain was recently generated by deleting four genes through CRISPR/Cas9-based editing [17]. These examples show the extraordinary potential of genetic strain engineering strategies based on the knowledge of cellulase gene regulation.

Notably, regulation of enzyme coding genes can vary substantially among related fungal species. The transcriptional activators for cellulolytic genes are ClrB in *A. niger* and *P. oxalicum* and Clr-2 in *Neurospora crassa* [24,25]. ClrA is another transcriptional regulator, whose deletion in *A. niger* had a minor effect on plant biomass deconstruction compared to the deletion of ClrB, while deletion of its orthologue Clr-1 in *N. crassa* led to strongly impaired cellulase production and severe growth defects on cellulose and cellobiose [26,27]. The transcription factor XlnR and its orthologues are involved in regulation of xylanolytic genes in *A. niger, P. oxalicum, T. reesei* and *N. crassa* [24,28,29]. In *A. niger*, XlnR is also involved in the activation of cellulolytic genes [2,27]. In *T. reesei*, the *xlnR* homolog *xyr-1* is the most important regulator of cellulases and xylanases, and its deletion leads to severe growth defects on cellulose [30]. These results make regulatory genes attractive targets for strain engineering purposes.

Development of genetic tools to improve the regulation of plant cell wall degrading enzymes in filamentous fungi provides a pathway to engineer a wider variety of hypersecreting fungal strains. The goal of this study was to (1) develop genetic tools, namely a transformation system, a CRISPR/Cas9 gene editing method and a sexual crossing protocol, for *T. aurantiacus* and (2) employ those tools to manipulate the xylanase regulator *xlnR* for a proof of principle study for strain engineering of xylanase secretion. Here we report on successful establishment of those objectives: an ATMT procedure was established, which was then used to implement the CRISPR/Cas9 system in *T. aurantiacus* by inactivating the native *pyrG* gene through mutations caused by the Cas9 endonuclease. Lastly, a sexual crossing protocol has been developed for this fungus, allowing rapid combination of genetic modifications within a week. As a proof of concept, we generated high xylanase secreting strains via integration of a *xlnR* cassette into the fungal genome with ATMT, displaying the applicability of the developed methods for generating high enzyme secreting *T. aurantiacus* strains for cost-efficient biofuel production.

## Results

### *Agrobacterium tumefaciens* mediated transformation system development for *T. aurantiacus*

Various transformation protocols have been developed for filamentous fungi, such as protoplast generation, electroporation, ATMT and nanoparticle-based methods such as biolistics [14]. Attempts to transform *T. aurantiacus* by protoplastation and electroporation were unsuccessful (data not shown). Therefore, ATMT was chosen for the transformation of T. *aurantiacus*.

*T. aurantiacus* is a homothallic fungus and was reported to only produce ascospores for propagation through self-crossing [31]; no conidiospores have been observed for this species. ATMT involves the co-cultivation of the bacteria with germinating spores of the fungus. We therefore first determined optimal culture conditions for *T. aurantiacus* ascospore production by testing the growth media PDA, Vogel’s minimal medium and YPD (data not shown). Spore production was found to be as follows: PDA > Vogel’s minimal medium > YPD. Since PDA yielded the largest number of spores, it was chosen for the following experiments. In the next step, we tested different pre-culture conditions for optimal spore production and germination rates. The most efficient spore germination was found when spores were harvested from PDA plates grown for 2 days at 50°C and 3 to 4 days at 45°C (Fig. 1a). However, a higher spore yield was obtained from plates, on which *T. aurantiacus* was grown for 4 days at 45°C (∼7 * 10^8^ spores per plate, see Fig. 1b). Therefore, the latter incubation time was chosen to harvest spores for ATMT.

**Fig. 1:**
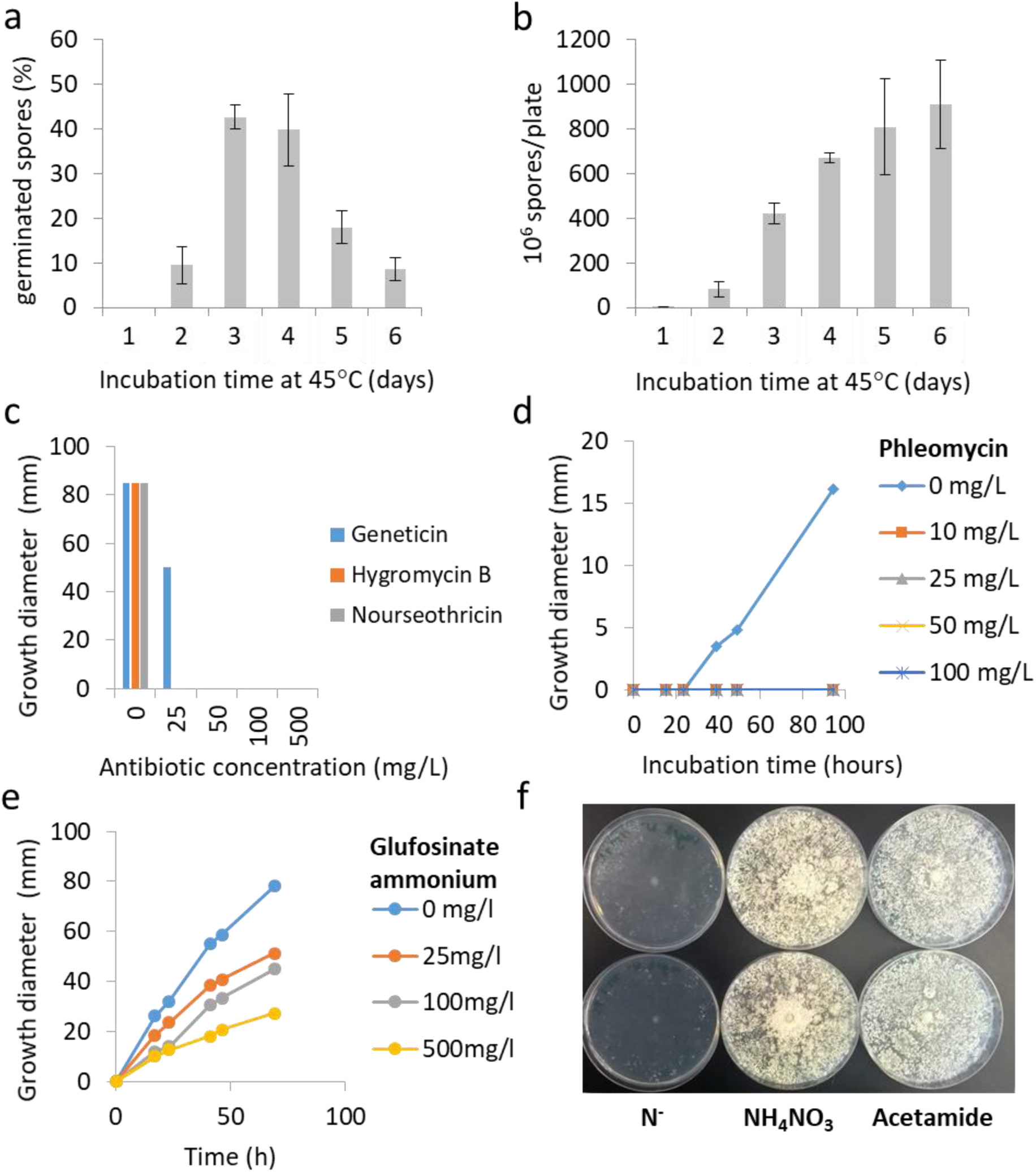
Ascospore production and antibiotic susceptibility of *T. aurantiacus*: (a) Germination rates were assessed from spores of fungal cultures incubated at 50°C for 2 days and then 45°C for 1-6 days. At the indicated time, spores were scraped from 3 replicate plates for each day; germination was detected via randomized counts of spore suspensions. (b) The total amount of produced spores was calculated with a hemocytometer. Growth tests of *T. aurantiacus* on different selection markers: (c) hygromycin B, nourseothricin and geneticin (PDA medium), (d) phleomycin (Vogel’s minimal medium) and (e) glufosinate ammonium (Vogel’s minimal medium). (f) *T. aurantiacus* is able to grow on acetamide (Vogel’s minimal medium with no nitrogen added, ammonia nitrate and acetamide from left to right). Two replicate plates were used for all assays. Note that all antibiotics or acetamide were separately sterile filtered and added to the media after autoclaving. Two biological replicates were used for each test.

Transformation of fungi usually involves the selection of transformants via antibiotic resistance markers [32]. Commonly used antibiotic resistance genes confer resistance to hygromycin B, nourseothricin, glufosinate ammonium, geneticin and phleomycin [14,33]. Alternatively, acetamide can be used as a nitrogen source to isolate successful transformants through integration of an acetamidase gene (*amdS*), since not all species possess this essential enzyme for acetamide utilization [34]. To test the potential application of these selection systems for *T. aurantiacus*, the basic resistance level of the wild type strain against the above-mentioned antibiotics was determined. In addition, we tested if the fungus can grow using acetamide as the sole nitrogen source. Strong growth inhibition was observed on plates containing hygromycin B, nourseothricin, geneticin and phleomycin, while the fungus was able to grow on Vogel’s medium supplied with glufosinate ammonium (Fig. 1c-e). *T. aurantiacus* showed robust growth on minimal media plates supplemented with acetamide as the sole nitrogen source (Fig. 1f), which was consistent with the presence of a putative *amdS* gene in the *T. aurantiacus* genome (https://mycocosm.jgi.doe.gov/Theau2/Theau2.home.html).

For the first approach to establish ATMT, the Golden Gate compatible plasmid pTS57 (Supplemental Table 1) was constructed to mediate ectopic integrations of genes of interest into the fungal genome and to allow selection using hygromycin B resistance (Fig. 2a). In the pTS57 plasmid, the *hph* gene is driven by the native *T. aurantiacus tef-1* promotor and there is a cloning site for genes of interest expressed by the native *T. aurantiacus gpd* promotor.

**Fig. 2:**
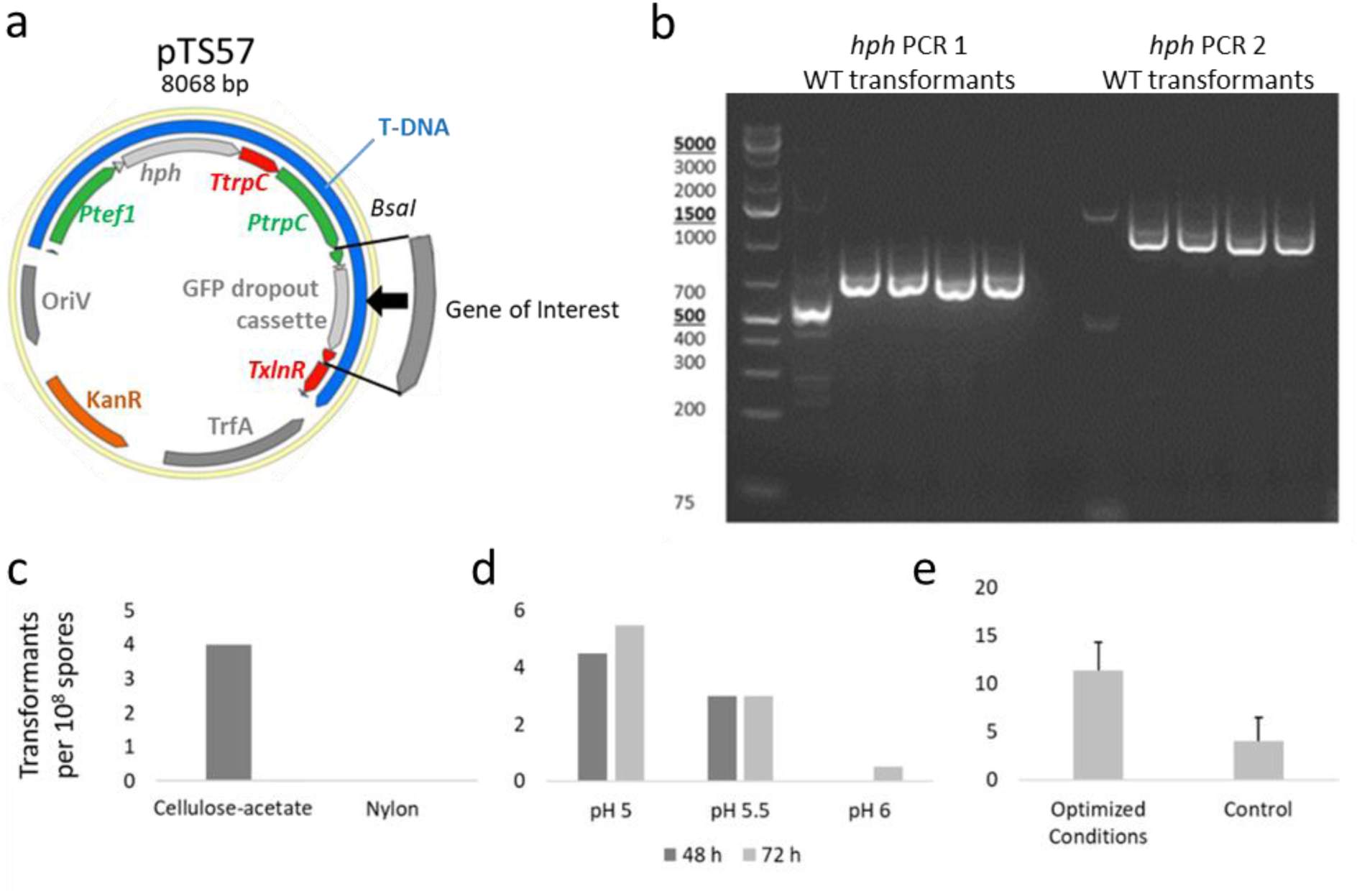
(a) The ATMT plasmid pTS57 was designed for efficient insertion of genes of interest and screening through Golden Gate Cloning upon replacement with a GFP-drop-out cassette. The gene of interest is expressed with the native *T. aurantiacus gpd*-promotor and *xlnR* terminator, the *hph* gene is expressed with the native *T. aurantiacus tef-1* promotor and *trpC* terminator. (b) PCR analysis to verify the *hph* integration into *T. aurantiacus* via ATMT. Optimization of the ATMT procedure for (c) membrane used, and (d) incubation time and pH. (e) A combination of optimized pH and temperature was tested regarding transformation rates. (c: 1 biological replicate, d: 2 biological replicates, and e: 3 biological replicates. Error bars indicate the standard deviation of 3 biological replicates). Two hygromycin B resistant strains are available through the JBEI Public Registry (Supplemental Table 2).

A previously developed ATMT protocol for *Rhodosporidium toruloides* [35] was modified for transformation of *T. aurantiacus* (for details see the Materials and Methods section). Briefly, 10^8^ fungal spores were mixed with 2 ml of an induced *A. tumefaciens* culture of OD_600_ of 1 carrying the plasmid pTS57 and incubated on a filter for 48 h on induction agar containing acetosyringone. After incubation, the spores were washed off the filters and spread on hygromycin B PDA containing cefotaxime to remove remaining bacteria. The grown fungal colonies were isolated after 2 days of incubation. In the initial experiment, four transformants were obtained from cellulose acetate filters using 10^8^ spores while no transformants were obtained when using a nylon filter (Fig. 2c). The presence of the hygromycin B resistance gene *hph* in all four strains was PCR verified through two different primer sets (Fig. 2b). Thus, this initial transformation approach was successful; however, transformation frequency was low.

The influence of the pH of the induction medium, the time of co-cultivation of the fungus and the bacteria, and the cultivation temperature was tested in order to further optimize the transformation protocol. We found that reducing the pH of the induction medium from 5.5 to 5 yielded on average 1-2 more transformants per 10^8^ spores, while varying the incubation time (48 vs 72 h) had virtually no effect on the number of transformants obtained (Fig. 2d). The temperature test indicated that increasing the temperature from 26°C to 28°C led to slightly higher transformation rates (data not shown). The combination of changing the induction medium pH to 5 and raising the incubation temperature to 28°C led to the isolation of ∼2.5 times more colonies compared to the initial conditions of pH 5.5 and 26°C (Fig. 2e).

### Genomic integration of *xlnR* expression cassettes lead to increased xylanase secretion

After establishing the ATMT procedure of *T. aurantiacus* ascospores, we used the method to demonstrate a proof of concept approach for the expression of a gene of interest in *T. aurantiacus*. Previous work had demonstrated that a continuous xylose feed induced both cellulase and xylanase activities in *T. aurantiacus*, raising the question of the involved transcriptional regulators [13]. In *T. reesei*, the transcription factor Xyr1 acts as an activator for xylanases and cellulases. This regulatory function is conserved for the respective homologs in different ascomycete species [30,36,37]. A *xyr1* homolog, named *xlnR*, had been identified in the *T. aurantiacus* genome in an earlier study [10]. To test the function of this regulator, we cloned the *xlnR* open reading frame into pTS67, where the gene is expressed by the native *T. aurantiacus gpd* promotor (Supplemental Table 1). The plasmid was transformed into the wild type reference strain by using the established ATMT protocol. 29 hygromycin B resistant transformants were obtained. For a subset of 16 isolates, the presence of the resistance gene within the genome was verified by PCR analysis using an *hph*-specific primer pair (data not shown). To test the effect of the newly integrated construct on xylanase activity, the 29 transformants and the wild type recipient strain were cultured in liquid media containing Avicel cellulose, a substrate that poorly induces xylanase, as the sole carbon source and xylanase activity was determined after 3 days of cultivation. For 24 out of the 29 isolates, a >50% increase in xylanase activity was observed, and 10 transformants out of this group demonstrated a >300% increase with one transformant displaying a 500% increase (Fig. 3a). The secretion of elevated amounts of xylanase by the transformants was also tested under non-inducing conditions. A shift experiment was performed with 4 isolates that displayed the highest amount of xylanase activity during incubations on Avicel celluose. These strains and the wild type were grown in glucose medium first and equal amounts of fungal biomass were then shifted to carbohydrate-free medium. We found a 6-fold increase in xylanase activity compared to wild type in these isolates (Fig. 3b). This proof of concept test indicated that enzyme secretion of this fungus could be successfully manipulated with the established ATMT procedure.

**Fig. 3:**
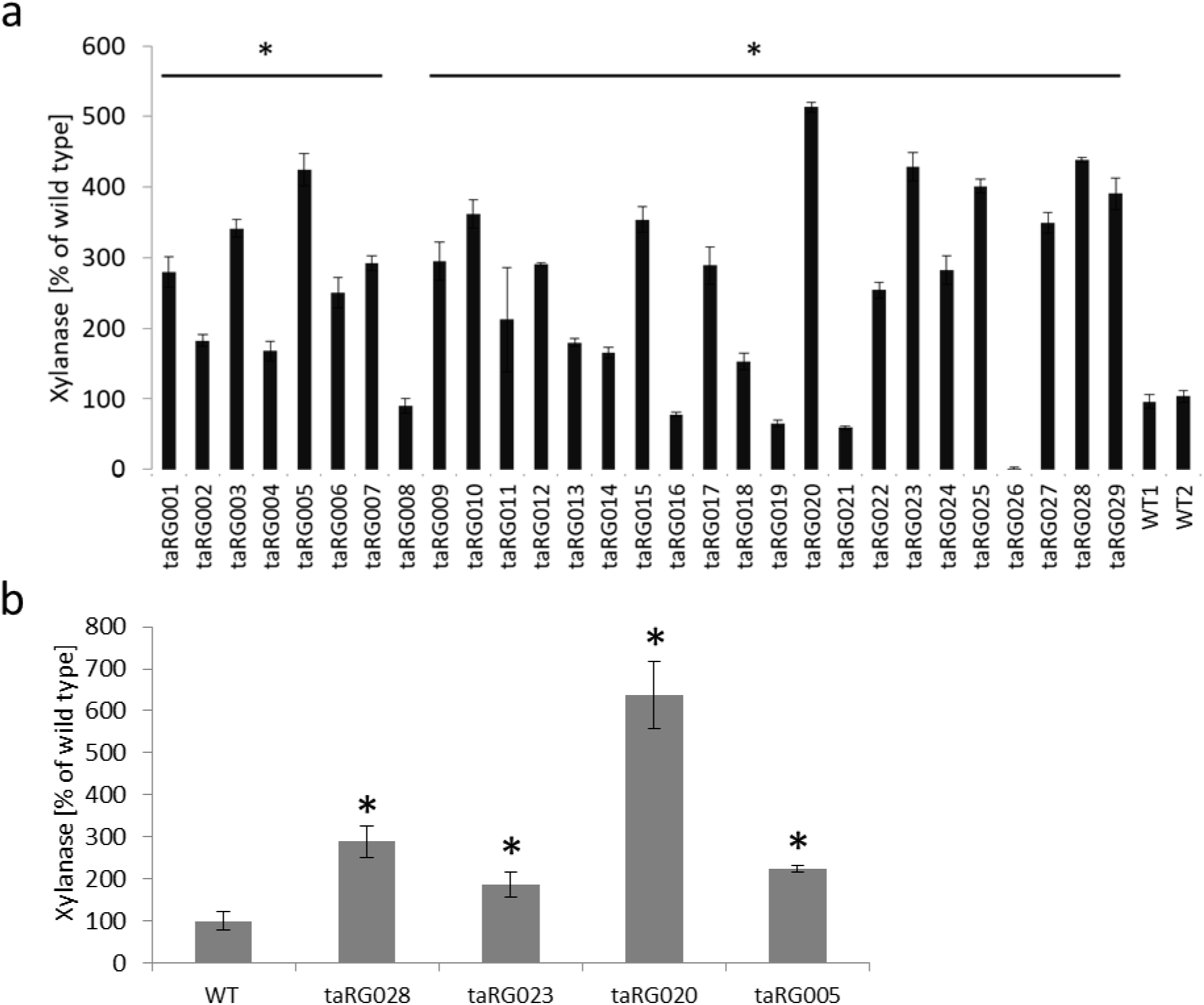
Xylanase activity of the *T. aurantiacus* strains transformed with a *Pgpd*::*xlnR* construct. (a) The DNS Assay was used to screen 29 transformants grown in Avicel medium for xylanase. (b) From a subset of the mutants tested in (a), a subset of 4 mutants displaying the highest xylanase activity was used for a shift experiment. The mutants were grown in McClendon’s medium supplemented with soy meal peptone and glucose for 48 hours and equal amounts of mycelium were shifted to starvation medium for 72 h and xylanase activity was measured. (a) bars represent one biological replicate and the error bars are the standard deviation of 3 technical replicates, the horizontal bars with the asterisk indicate statistical significant difference to the wild type strains (pval < 0.05), (B) bars and standard deviation are derived from three biological replicates, the asterisk indicate statistical significant difference to the wild type strains (pval < 0.05). The 10 *xlnR*-integrated strains with >300% increase in xylanase activity in Avicel-grown cells are available from the JBEI Public Registry (Supplemental Table 2).

### Development of a sexual crossing protocol for *T. aurantiacus*

*T. aurantiacus* is a self-fertile, homothallic fungus, which completes its sexual life cycle without a crossing partner. However, other homothallic species, such as the model fungus *Sordaria macrospora*, are often able to outcross [38]. In these cases, the basis for outcrossing is the formation of heterokaryotic mycelia via vegetative hyphal fusion of genetically compatible strains. Completion of the sexual cycle of these heterokaryons gives rise to genetically recombinant progeny. In order to test if outcrossing occurs in *T. aurantiacus* and to establish a crossing protocol, two strains with different selectable markers were employed. The hygromycin B resistant *T. aurantiacus* strain taRG008 (Fig. 3a) carrying the *xlnR* expression cassette described above was chosen as one of the crossing partners. For the second crossing partner, UV mutagenesis of *T. aurantiacus* ascospores was performed to isolate mutants that were uracil auxotrophs and resistant to 5-fluoroorotic acid (5-FOA). Metabolism of 5-FOA by wild type fungi generates the toxic intermediate fluorodeoxyuridine. 5-FOA therefore selects for mutants with non-functional *pyrG*, which encodes for orotidine 5’-phosphate decarboxylase and *pyrE*, which encodes for orotate phosphoribosyltransferase [39,40]. UV mutagenesis yielded two 5-FOA resistant strains (FOAR1 and FOAR2) that were isolated on 5-FOA minimal medium plates containing uracil. Subsequent sequencing of the *pyrE* gene region identified causative mutations for 5-FOA resistance (Fig. 4a). An insertion of 190 bp was found in FOAR1, which turned out to be a duplication of a part in the *pyrE* gene sequence while FOAR2 had a 1 bp insertion in *pyrE*, which created a frameshift mutation for both strains. FOAR2 was chosen as the partner to be crossed with the hygromycin B resistant strain taRG008 (Fig. 3a). Recombinant progeny were expected to harbor both resistances that could be easily screened for on media supplemented with hygromycin B, uracil and 5-FOA. The plate set-up for fungal crossings is shown in Supplemental Figure 1. Briefly, 2 fungal strains were inoculated on a PDA plate supplemented with uracil in alternating fashion to maximize the possibility to form a contact interface. From this interface that was expected to contain the crossed spores of both strains, the mycelium was scraped off the surface with a spatula and eluted in water. The spores were released through vortexing and filtered. Different dilutions were made and spread onto squared agar plates containing hygromycin B, 5-FOA and uracil to yield only very few (< 10) growing colonies, which simplified the isolation. Six progeny colonies (P1-6) were randomly isolated for further analysis on the selective plates. Genomic DNA was extracted from those colonies and was used to verify the integration of the *hph* gene cassette that was passed on from parent strain taRG008 (Fig. 4b, left gel) as well as the *pyrE* mutation of the parent strain FOAR2 (Fig. 4c). The wild type, FOAR1, FOAR2, the *xlnR*/*hph* expressing strain taRG008 and one progeny isolate (P1) were included as controls for both PCRs (Fig. 4b, right gel). The PCR amplification of the *hph* gene and Sanger sequencing of the *pyrE* PCR confirmed that both modifications were only present in the progeny isolates (Fig. 4c).

**Fig. 4:**
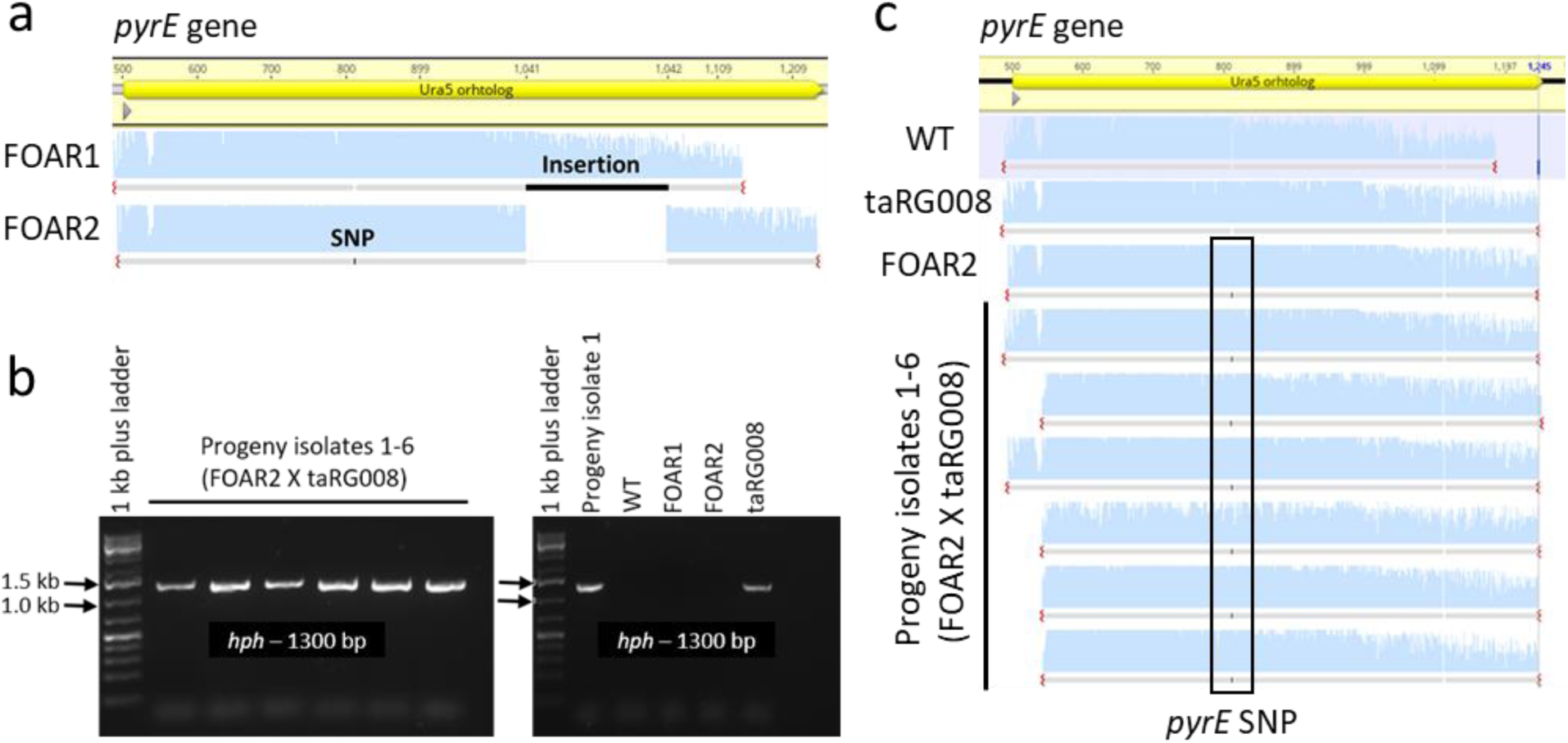
Testing sexual outcrossing in *T. aurantiacus*: (a) Sequencing data of the *pyrE* gene of two 5-FOA resistant isolates (FOAR1-2) were aligned to the *pyrE* reference sequence (primers are listed in Table 2). FOAR2 was crossed with the hygromycin B resistant strain taRG008 (*hph* strain). The progeny of this cross was analyzed through (b) PCR amplification of the *hph* gene (wild type, FOAR strains and taRG008 were included as controls) (c) Sequencing data of the *pyrE* gene sequence of the crossed strains in (b) were aligned to the native *pyrE* reference sequence. The sequence analysis was performed with Geneious version 11.1 (Biomatters). This analysis indicated that only the progeny isolates displayed genomic integrations of the *hph* gene and the *pyrE* mutation from FOAR2. FOAR1-2 and an FOAR2 *x* taRG008 crossed strain are available from the JBEI Public Registry (Supplemental Table 2).

### Development of a CRISPR/Cas9 protocol for gene deletion in *T. aurantiacus*

CRIPSR/Cas9 is a powerful genome editing tool consisting of an RNA-guided endonuclease (Cas9) and one or multiple guide RNAs (gRNAs) for targeting one or several genomic loci at the same time [41]. Cas9 can be introduced into the fungal cell in the form of DNA, RNA or a protein-RNA complex. Unlike other commonly used transformation strategies, ATMT only allows transformation of DNA fragments into the fungal cell. Therefore, ATMT-mediated Cas9 introduction into fungal cells relies on genomic integration of the Cas9 and gRNA expression cassettes [42,43]. To apply Cas9-based editing in *T. aurantiacus*, an AMA1 plasmid-based expression approach from Nødvig [44] was chosen and modified for ATMT. In the present study, Cas9 and gRNA expression cassettes were amplified from AMA1 based Cas9 plasmids generated in the study mentioned before and integrated into the ATMT plasmid pTS57, generating a new series of plasmids (pJP1, pJP3, Table 1). These ATMT compatible Cas9 plasmids were then used to integrate an expression cassette of the Cas9 gene, the gRNA and the *hph* marker into the fungal genome.

To demonstrate the CRISPR/Cas9 gene editing approach in *T. aurantiacus*, the *pyrG* gene was chosen as a target for gene inactivation. Disruption of *pyrG* causes uracil auxotrophy but also confers resistance to 5-flouroorotic acid (5-FOA), which allows screening of *pyrG* mutants. Three different gRNAs targeting the *pyrG* gene in *T. aurantiacus* were designed using CRISPOR web-tool [36] (Supplemental Table 4) and first tested *in vitro* through performing a Cas9 cleavage assay. Accordingly, purified Cas9 protein, a *pyrG* PCR product and one *in vitro* transcribed gRNA per reaction were incubated to facilitate *pyrG*-DNA cleavage mediated by the Cas9 ribonucleoprotein complex. Each reaction was then analyzed through agarose gel electrophoresis to visualize the cleaved DNA fragments. The cleavage efficiency was found to be as follows: gRNA 1 > gRNA 3 > gRNA 2 (Fig. 5a). Therefore, gRNA 1, gRNA 3 and the Cas9 gene were cloned into the ATMT plasmid pTS57, yielding the plasmids pJP1 and pJP3, respectively. Transformations of *T. aurantiacus* ascospores using those plasmids were performed by ATMT. The selection for positive transformants was performed through screening for hygromycin B resistance (Fig. 5b) and, on average, approximately 5.5 (pJP1) and 3.5 (pJP3) transformants per 10^8^ spores were obtained. A subset of 20 of each of these transformants were randomly picked and further screened for 5-FOA resistance on Vogel’s MM supplemented with uracil and 5-FOA. Three (pJP1) and 12 (pJP3) out of 20 transformed *T. aurantiacus* isolates displayed 5-FOA resistance (Supplemental Fig. 2a, Fig. 5c). Sequencing of a subset of isolates from transformations with pJP1 and pJP3 revealed mutations within the protospacer targeting sequence of the *pyrG* gene, confirming that Cas9 cleavage led to base deletions and mismatches next to the PAM sequence, which caused a frameshift in the *pyrG* gene in all sequenced strains and thus 5-FOA resistance of the respective strains (Supplemental Fig. 2b). The deletion efficiency was then calculated based on the fraction of *T. aurantiacus* transformants isolated on hygromycin B medium after the ATMT transformation that also had a mutation in the *pyrG* gene: gRNA 1 displayed a deletion efficiency of 10% while gRNA 3 displayed a deletion efficiency of 35% (Fig. 5c). Thus, the Cas9 system successfully introduced mutations in the *pyrG* gene, and selection on 5-FOA turned out to be effective to screen for those mutants.

**Fig. 5:**
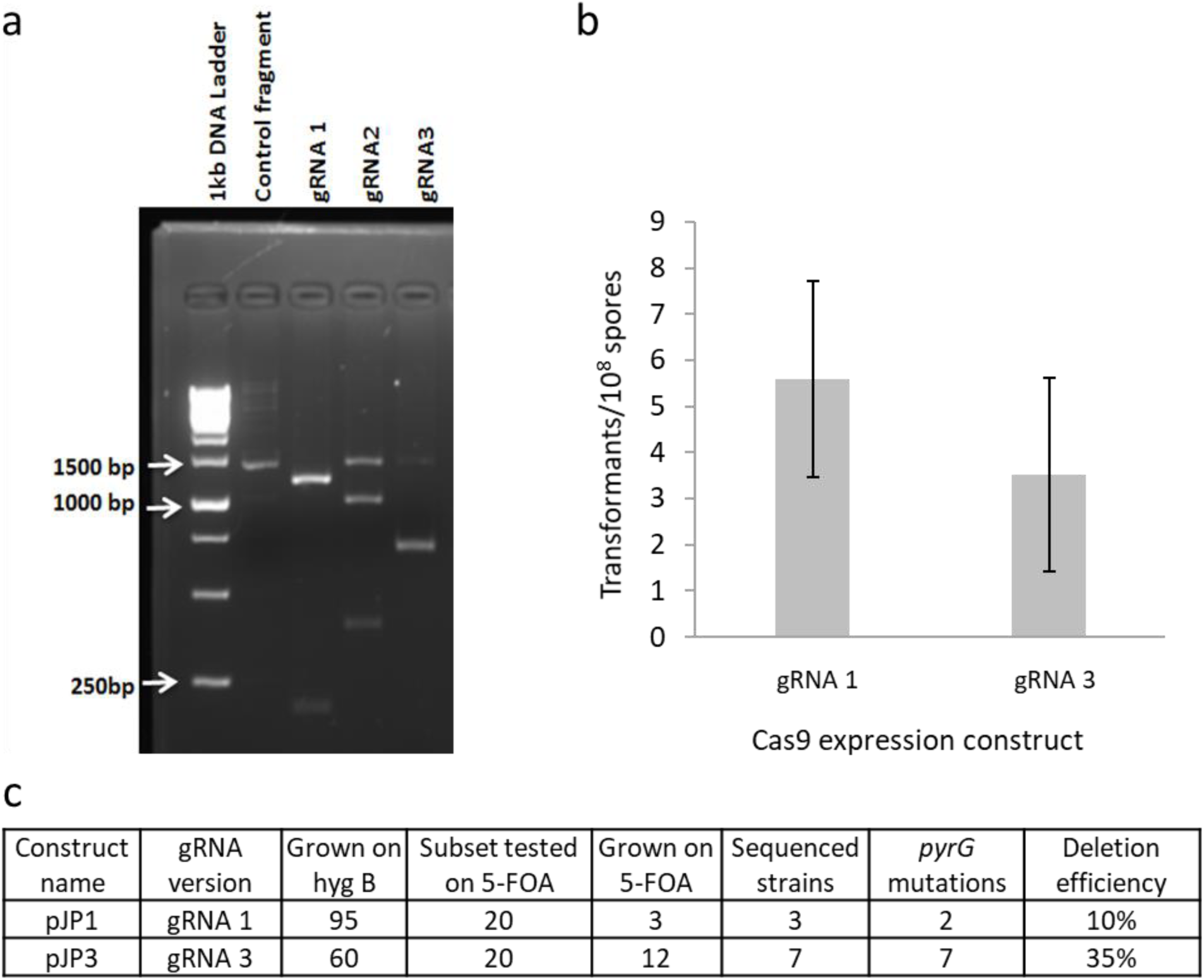
CRISPR/Cas9 development in *T. aurantiacus*: (a) *in vitro* Cas9 cleavage assay: Agarose gel depicting the uncleaved control fragment and the Cas9 cleavage of the target *pyrG* sequence with gRNA 1, 2 and 3. (b) Transformation efficiency per 10^8^ spores with gRNA 1 and 3 containing vectors, selected on hygromycin B uracil plates. Each bar displays the mean and standard deviations from 17 biological replicates. (c) Deletion efficiencies of both gRNAs targeting Cas9 to the *pyrG* gene in *T. aurantiacus*. Six *T. aurantiacus* strains transformed with pJP1 are available from the JBEI Public Registry (Supplemental Table 2).

## Discussion

In this study, we have established a variety of genetic tools to engineer *T. aurantiacus*. These tools include an ATMT method for transformation, a sexual crossing protocol and a Cas9-based method for gene editing. While genetic tools have been developed for a number of mesophilic filamentous fungi, there are limited genetic tools for thermophilic fungi, so development of genetic tools for *T. aurantiacus* represents the first step towards establishing this fungus as a production platform for thermostable enzymes.

ATMT was a successful approach to transform *T. aurantiacus*; however, the process is more time-consuming than other frequently used transformation approaches and limits the extent of engineering possibilities. Developing protoplast transformation or electroporation protocols for *T. aurantiacus* will accelerate and expand engineering. ATMT was previously established for conidiospores of the thermophilic fungus *M. thermophila* [16] and generated up to 145 transformants per 10^5^ spores. Thus, the transformation rates reached for *T. aurantiacus* ascospores in this study (10 per 10^8^ spores) were significantly lower. The ATMT procedure was then used to genomically integrate an expression cassette of the transcriptional regulator *xlnR* (Theau_38177). Transformants carrying the *xlnR* construct exhibited high variability of xylanase activity in the culture supernatants, which would be consistent with random integrations of the cassettes in unknown genomic regions and variable numbers of genetic copies inserted into the genome. Nevertheless, up to 500% increased xylanase activity was observed compared to the wild type in strains carrying the *xlnR* construct. Therefore, the *T. aurantiacus xlnR* appears to have a comparable function to its homologs in the closely related *Aspergillus spp*. and *P. oxalicum*, which regulate xylanase gene expression [24,27,37]. Notably, cellulases and xylanases secretion of *A. niger* in the presence of D-xylose was linked to phosphorylation of XlnR, which mediates the induction of the respective genes in the presence of this carbon source [2,29]. *T. aurantiacus* is closely related to *A. niger* and was found to produce high amounts of cellulases and xylanases during D-xylose fed-batch conditions, which might be mediated by XlnR as well [13].

Furthermore, the ATMT method enabled the successful establishment of the CRIPSR/Cas9 system in *T. aurantiacus*. The gene editing system relied on the non-homologous end joining (NHEJ) repair pathway to generate mutations in the *pyrG* gene, which has been previously demonstrated in *M. thermophila* for the *amdS* gene [17]. Additionally, the ability to generate protoplasts for *M. thermophila* led to the introduction of multiple plasmids, which permitted deletions of genes using homology-directed repair (HDR) mechanisms with a *ku70* deletion strain [16]. Nonetheless, the CRISPR/Cas9 system can now be used to modify and investigate the role of other well-known regulators, such as *creA, clrA, clbR* and *amyR* in a multiplexed manner to further uncover cellulase and xylanase regulation in *T. aurantiacus* [27,33,37,45,46]. Moreover, other genes related to carbon catabolite repression and secretion of other carbohydrate active enzymes might be vital targets for understanding and engineering CAZyme secretion in *T. aurantiacus* [47,48]. Finally, recyclable markers such as *pyrG* allow to delete target genes with a high efficiency and then remove the marker through a loop-out mechanism by adding homology repeats [49].

The demonstration of sexual crossing between two strains of *T. aurantiacus* reveals an important advantage for this fungus as a potential platform for producing thermostable enzymes. Crossing under laboratory conditions is a valuable genetic tool only available for a limited number of species, such as the model fungi *N. crassa* [50], *A. nidulans* [51] and recently also *T. reesei* [21], but is lacking for several industrially-relevant fungi with unknown teleomorphs such as *A. niger* and *A. oryzae* [20,22]. *M. thermophila* is not capable of self-crossing and does not cross with close relative *Myceliophthora heterothallica*, which has been experimentally demonstrated to have a sexual life cycle [52,53]. An additional advantage of the homothallic *T. aurantiacus* is that crossing does not require strains with different mating types as in heterothallic fungi like *T. reesei* [54] or *P. chrysogenum* [55]. Ascospores are the only means of propagation in *T. aurantiacus* and are produced in as little as 4-5 days. Since ascospores originate from a single nucleus, the resulting progeny are always homokaryotic, allowing for quick and simple purification of originally heterokaryotic transformants. Additionally, the sexual crossing of *T. aurantiacus* was demonstrated on conventional fungal media such as PDA. Since crossed transformants were isolated within a week, it appears that the crossing procedure with this fungus is substantially faster and easier than procedures used for other fungi such as *N. crassa* or *A. nidulans* [56].

In summary, the developments demonstrated in this paper will enable rapid stacking of genetic modifications into new strains for subsequent strain tests. We expect these developments and further improvements of the genetic transformation procedure to turn *T. aurantiacus* into a novel host for studying plant cell wall deconstruction, sexual biology and cell biology. In addition, these protocols provide the basis for developing *T. aurantiacus* as a host for numerous biotechnological applications.

## Conclusion

The methods generated in this study will enable to substantially expand the use of *T. aurantiacus* in both applied and fundamental studies. *T. aurantiacus* is an intriguing host for cellulase production due to the extraordinary thermostability of its cellulases, the high enzyme titers secreted by the wild type and, since it is a homothallic fungus, the possibility to rapidly cross strains carrying different mutations into homokaryotic progeny in substantially shorter time frames than currently used industrial fungi, thereby enhancing strain engineering. With further development regarding the transformation system, CRISPR/Cas9, and the crossing protocol, it will be possible to generate genetically modified strains that can be crossed to combine desired mutations. This will enable high CAZyme production with *T. aurantiacus* through deleting or overexpressing regulators and other genes known to impact CAZyme production in related filamentous fungi.

## Methods

### Chemicals

All chemicals were purchased from Sigma-Aldrich unless otherwise indicated.

### Strains and culture conditions

*T. aurantiacus* ATCC® 26904™ was obtained from the American Type Culture Collection and grown on TEKNOVA potato dextrose agar (PDA) plates to obtain ascospores for transformation purposes. The PDA plates were inoculated with ascospores and incubated for two days at 50 °C before they were transferred to 45 °C for another four days. This shift was performed due to elevated evaporation of PDA plates at 50 °C. The plates were covered with a glass beaker to reduce drying, and plastic containers filled with distilled H_2_O provided a moist atmosphere. Cultivation of the uracil auxotroph strains generated in this study was performed on solid Vogel’s minimal medium containing Vogel’s salts solution, 2 % sucrose and 1.5 % bacto agar supplemented with 1 g/L uracil and 1 g/L 5-FOA as indicated.

*Agrobacterium tumefaciens* strain EHA105 was grown Luria-Bertani (LB) medium plates (supplemented with kanamycin at 50 µg ml^-1^ when culturing transformed strains harboring plasmids for the fungal transformations). After two days, 2-3 *A. tumefaciens* colonies carrying the desired plasmids were inoculated in 10 ml of liquid LB medium at 30°C supplemented with kanamycin as described above.

### Antibiotic resistance plate tests of *T. aurantiacus*

For all plate tests, counted *T. aurantiacus* spores were placed in the center of 9 mm agar plates containing the desired antibiotic. These plates were then incubated at 45°C, and fungal growth was measured after 72 h (hygromycin B, geneticin, nourseothricin, 5-fluoroortic acid [5-FOA] and 5-fluoroacetamide [5-FAA]) or as indicated (glufosinate ammonium and phleomycin). The fungal mycelium diameter was measured with a vernier caliper from two sides and averaged. Antibiotic concentrations were added as indicated or 1.3 mg/ml for 5-FOA and 5-FAA. All antibiotics were sterile filtered separately and added after sufficient cooling of the agar. The following media compositions were used: PDA for hygromycin B, geneticin and nourseothricin, Vogel’s minimal medium with 2% sucrose for glufosinate ammonium, phleomycin (pH 8), 5-FOA and 5-FAA.

### Ascospore production and germination rate tests

PDA plates (Sigma-1879V) were inoculated in 3 biological replicates for each time point and ascospores were incubated at 50°C for 2 days and 1 – 6-das at 45°C. Spores were harvested through scraping off the surface with a cell spreader tow times and filtering through miracloth. These spores were counted with a hemocytometer, diluted appropriately, spread on a new plate and incubated for 16 hours at 45°C. These spores were then randomly imaged with a Leica-DM4000B microscope, and the germination rate was calculated via counting germinated versus non germinated spores with ImageJ [57]. A minimum of 385 spores was counted for each day, except for day 3, where almost no spores were present.

### Plasmid design and cloning strategy

The base vector pTS57 (P*gpd*::P::*gfp*::T::T*xlnR*; P*tef1*::*hph*::T*trpC*) was used to generate all further vectors (see Supplemental Table 1). pTS57 expresses the gene of interest with the *T. aurantiacus gpd* promotor and *xlnR* terminator, which flank a *gfp*-dropout cassette that is recognized by *E. coli*. This cassette has two *BsaI* restriction sites at either end and allows genes of interest to be inserted through Golden Gate Cloning. *E. coli* transformants harboring the plasmid with the integrated gene of interest can then be identified through loss of *gfp* fluorescence on a blue-screen. Additionally, pTS57 contains the hygromycin B phosphotransferase (*hph*) expressed by the native *T. aurantiacus tef-1* promotor and *trpC* terminator. Plasmids were isolated after assembly and electroporation into MEGAX DH10B T1R Electrocomp Cells (Thermo Fisher Scientific) with the QIAprep Spin Miniprep Kit (Qiagen) and transformed into *A. tumefaciens* strain EHA105 through electroporation. The plasmid pTS67 (see Supplemental Table 1) was also used, which was derived from pTS57 by the above-mentioned procedure to constitutively express the transcription factor *xlnR*.

ATMT compatible CRISPR/Cas9 plasmids were designed to target the *pyrG* gene in the target host *T. aurantiacus*. The target sequences were obtained from JGI mycocoms. All plasmid maps were designed using the software Geneious 11.1.2 (https://www.geneious.com). The gRNAs used in this study were designed using the CRISPOR algorithm [58] (http://crispor.tefor.net) to obtain predicted guide sequences for PAMs in the target gene. Three different gRNA sequences (protospacers) with no predicted off-targets were chosen and tested *in vitro* for correct cleavage of the target sequence by Cas9 endonuclease before performing *in vivo* transformation experiments. [44]

All steps for the gRNA synthesis were followed according to the GeneArt Precision gRNA Synthesis Kit (Thermo Fisher Scientific, Waltham, MA, United States). Then, the *in vitro* Cas9 cleavage was performed using a previously amplified target *pyrG* amplicon following the steps of the Guide-it™ sgRNA In Vitro Transcription and Screening Systems User Manual (Takara Bio USA, Inc., Mountain View, CA, United States).

The two gRNAs with the highest cleavage efficiency (gRNA1 & 3) were inserted into vector pFC334 and then combined with the Cas9 gene from pFC332 via USER Cloning as described by Nødvig et al. [44]. The resulting plasmids were named JP36_1 (gRNA1) and JP36_3 (gRNA3). Those plasmids were then used as templates to insert their gRNA-cas9-expression-cassette into the ATMT vector pTS57 through Gibson Assembly yielding. the ATMT compatible Cas9 plasmids pJP1 (gRNA1) and pJP3 (gRNA3).

### ATMT transformation procedure

*T. aurantiacus* ascospore preparation and *A. tumefaciens* cultivation were performed as described above. The solid and liquid *A. tumefaciens* induction medium contained 200 µM acetosyringone (induction medium: salts, phosphor buffer, MES-buffer, glucose, thiamine, acetosyringone and water, see [35]). All reagents were sterile filtered with Corning filter systems or small filters and a sterile syringe. The pH of the induction medium was adjusted to pH = 5.

A modified version of the ATMT procedure for *Rhodosporidium toluroides* was used [35]. Briefly, ascospores of *T. aurantiacus* were harvested from 6-day old PDA plates and counted with a hemocytometer. *A. tumefaciens* EHA105 was grown overnight in 10 ml of liquid LB medium containing 50 µg/ml kanamycin. From this culture, a new liquid LB-kanamycin culture was generated with optical density at 600 nm (OD_600_) of 0.5 that was grown to OD_600_=1 and then pelleted, washed three times with induction medium, resuspended in induction medium and incubated for further 24 hours. Freshly harvested fungal spores and *A. tumefaciens* cell cultivated in induction medium overnight were filtered onto a 0.45 µm cellulose acetate membrane (0.45 µm MCE Membrane, MF-Millipore) and incubated on induction medium agar plates for 2 days. The spores and cells were washed off with a wash solution containing 200 µg/ml of cefotaxime and were spread on PDA plates containing 200 µg/ml of cefotaxime and 50 µg/ml of hygromycin B with subsequent incubation for 3 days at 45 °C. Colonies were isolated and grown on a fresh PDA plates containing 200 µg/ml of cefotaxime and 50 µg/ml of hygromycin B to remove untransformed spores through harvesting ascospores from the proximate region for generating cryostocks and performing further strain tests. For Cas9 tests, those colonies were then screened for 5-FOA resistance due to CRISPR-mediated mutations in *pyrG* on Vogel’s minimal medium containing 2% sucrose, 1 mg/ml 5-FOA and 1 mg/ml uracil.

### Strain tests and screening of transformants

For cellulase and xylanase activity tests, strains isolated from hygromycin B PDA plates were used to inoculate McClendon’s medium,0.8% SMP and a carbon source as indicated (Avicel cellulose or no carbon added). For enzyme assays, 0.8 ml of the culture broth was filtered through a spin filter column (Mini Spin Column, EconoSpin). The enzyme assays were performed on a Biomek FX through a DNS method. The first step involved manually adding 75 µl of 1% w/v Beechwood xylan (Megazyme) solution to a 96 well PCR plate (FLAT 96 WELL PCR PLATE, VWR) and 5 µl of enzyme solution. The Biomek FX was used to add DNS reagent to the PCR plates. Upon incubation of these plates at 95 °C, the plate content was transferred with Biomek FX to a flat bottom 96 well plates, and the absorbance was measured at 540 nm. D-glucose was used as a standard for the CMCase assay and D-xylose for the xylanase assay. Uracil auxotrophic strains were isolated on 5-FOA agar as described above and inoculated in PD broth containing 1 g/L uracil.

For strain verification, the mycelium DNA was extracted with the Maxwell RSC Plant DNA Kit (Promega) on the Maxwell RSC Instrument (Promega) according to the manual. One modification involved bead beating of intact mycelium with 300 µl extraction buffer. The concentration of the isolated DNA was measured with NanoDrop 2000 and used for PCR verifications of successful transformation. All sequencing verification was performed through Sanger sequencing.

### Sexual crossings

The mutant *T. aurantiacus* strains taRG008 (hygromycin B resistant) and FOAR2 (5-FOA resistant) were first grown individually as described above. A PDA-uracil petri dish was divided in four quarters and spore suspensions from taRG008 and FOAR2 were spotted on the middle of each quarter in an alternating fashion (S. Fig. 1). Incubation was performed at 45 °C for six days. Once a lawn of ascospores was produced, spores were scraped off at the interface of the two crossing strains with a sterile spatula, transferred into 750 μL sterile H_2_O, vortexed, and filtered through a sterile filter tip with miracloth. A dilution was prepared and plated onto a 12×12 cm square plate with Vogel’s minimal medium supplemented with 1 g/L 5-FOA, 1 g/L uracil and 50 µg/mL hygromycin B in triplicates. After incubation at 45 °C for three days, growth was visible. Randomly picked colonies were isolated and grown on the same media as the isolation plates. Genomic DNA was extracted from isolated colonies and used for PCR-based verification purposes.

## Authors’ contributions

RG, SWS, AF and TS designed experiments; RG conducted experimental antibiotic concentration and spore germination tests; SH, JP, CRV, AO, LLK, RG and TS designed and constructed the ATMT plasmids; RG, MJ, PC, SH, AO, LM and JG performed experimental work related to ATMT establishment; JP, CRV, PC, RG, LLK, SF and LF performed the experimental CRISPR/Cas9 establishment. JP, RG, CRV, PC and LF performed the sexual crossing protocol establishment; RG, JP, and MJ performed data analysis; RG, SWS and AF wrote the manuscript. All authors read and approved the final manuscript.

## Acknowledgements

This work was supported by the German Academic Exchange Service (DAAD) through a granted stipend to R.G. (Jahresstipendien für Doktorandinnen und Doktoranden, Studienjahr 2018/19 under the program number 57380837). Additional funding for student internships was provided by the Austrian Marshall Plan Foundation (J.P., M.J., L.F., R.G.). Portions of this work were funded by the U.S. Department of Energy, Office of Energy Efficiency and Renewable Energy, Bioenergy Technologies Office. Portions of this work were performed as part of the DOE Joint BioEnergy Institute (http://www.jbei.org) supported by the U.S. Department of Energy, Office of Science, Office of Biological and Environmental Research, through contract DE-AC02-05CH11231 between Lawrence Berkeley National Laboratory and the U.S. Department of Energy. Finally, we want to acknowledge the help of Dai Zyu and Jon Magnuson from the Pacific Northwest National Laboratory in consulting and sharing resources related to ATMT development in *T. aurantiacus*. U.S. Government retains and the publisher, by accepting the article for publication, acknowledges that the U.S. Government retains a nonexclusive, paid up, irrevocable, worldwide license to publish or reproduce the published form of this work, or allow others to do so, for U.S. Government purposes.

## References

1. Popper ZA, Michel G, Herve C, Domozych DS, Willats WGT, Tuohy MG, et al. Evolution and Diversity of Plant Cell Walls□: From Algae to Flowering Plants. Annu Rev Plant Biol. 2011;62:567–90.

2. Glass NL, Schmoll M, Cate JHD, Coradetti S. Plant Cell Wall Deconstruction by Ascomycete Fungi. Annu Rev of Microbiology. 2013;477–98.

3. Luo Y, Lee J, Zhao H. Challenges and opportunities in synthetic biology for chemical engineers. Chem Eng Sci. Elsevier; 2013;103:115–9.

4. Zhao Z, Liu H, Wang C, Xu J. Comparative analysis of fungal genomes reveals different plant cell wall degrading capacity in fungi. BMC Genomics. BMC Genomics; 2013;14:1.

5. Tian C, Beeson WT, Iavarone AT, Sun J, Marletta M a, Cate JHD, et al. Systems analysis of plant cell wall degradation by the model filamentous fungus *Neurospora crassa*. Proc Natl Acad Sci U S A. 2009;106:22157–62.

6. Peterson R, Nevalainen H. *Trichoderma reesei* RUT-C30 – Thirty years of strain improvement. Microbiology. 2012;158:58–68.

7. Visser H, Joosten V, Punt P, Gusakov A, Olson P, Joosten R, et al. Development of a mature fungal technology and production platform for industrial enzymes based on a *Myceliophthora thermophila* isolate, previously known as *Chrysosporium lucknowense* C1. Ind Biotechnol. 2011;7(3):214–23.

8. Blumer-Schuette SE, Brown SD, Sander KB, Bayer EA, Kataeva I, Zurawski J V., et al. Thermophilic lignocellulose deconstruction. FEMS Microbiol Rev. 2014;38:393–448.

9. Kolinko S, Wu YW, Tachea F, Denzel E, Hiras J, Gabriel R, et al. A bacterial pioneer produces cellulase complexes that persist through community succession. Nat Microbiol. 2018;3:99–107.

10. Schuerg T, Gabriel R, Baecker N, Baker SE, Singer SW. *Thermoascus aurantiacus* is an Intriguing Host for the Industrial Production of Cellulases. Curr Biotechnol. 2016;6:89–97.

11. Berka RM, Grigoriev I V, Otillar R, Salamov A, Grimwood J, Reid I, et al. Comparative genomic analysis of the thermophilic biomass-degrading fungi *Myceliophthora thermophila* and *Thielavia terrestris*. Nat Biotechnol. 2011;29:922–7.

12. McClendon SD, Batth T, Petzold CJ, Adams PD, Simmons BA, Singer SW. *Thermoascus aurantiacus* is a promising source of enzymes for biomass deconstruction under thermophilic conditions. Biotechnol Biofuels. 2012;5:54.

13. Schuerg T, Prahl JP, Gabriel R, Harth S, Tachea F, Chen CS, et al. Xylose induces cellulase production in *Thermoascus aurantiacus*. Biotechnol Biofuels. 2017;10:1–11.

14. Meyer V. Genetic engineering of filamentous fungi – Progress, obstacles and future trends. Biotechnol Adv. 2008;26:177–85.

15. Bevan M. Binary Agrobacterium vectors for plant transformation. Nucleic Acids Res. 1984;12:8711–21.

16. Xu J, Li J, Lin L, Liu Q, Sun W, Huang B, et al. Development of genetic tools for *Myceliophthora thermophila*. BMC Biotechnol. 2015;15:1–10.

17. Liu Q, Gao R, Li J, Lin L, Zhao J, Sun W, et al. Development of a genome-editing CRISPR/Cas9 system in thermophilic fungal *Myceliophthora* species and its application to hyper-cellulase production strain engineering. Biotechnol Biofuels. 2017;10:1.

18. Wheeler BHE. Genetics of Fungi. Annu Rev Microiol. 1957;12:365–82.

19. Ebbole D, Sachs MS. A rapid and simple method for isolation of *Neurospora crassa* homokaryons using microconidia. Fungal Genet Rep. 1990;37:7.

20. Dyer PS, O’Gorman CM. Sexual development and cryptic sexuality in fungi: insights from *Aspergillus* species. FEMS Microbiol Rev. 2012;36:165–192.

21. Seidl V, Seibel C, Kubicek CP, Schmoll M. Sexual development in the industrial workhorse *Trichoderma reesei*. Proc Natl Acad Sci. 2009;106:13909–14.

22. Kwon-Chung K, Sugui J. Sexual reproduction in *Aspergillus* species of medical or economical importance: Why so fastidious? Trends Microbiol. 2010;17:481–7.

23. Yao G, Li Z, Gao L, Wu R, Kan Q, Liu G, et al. Redesigning the regulatory pathway to enhance cellulase production in *Penicillium oxalicum*. Biotechnol Biofuels. 2015;8:71.

24. Gao L, Li Z, Xia C, Qu Y, Liu M, Yang P, et al. Combining manipulation of transcription factors and overexpression of the target genes to enhance lignocellulolytic enzyme production in *Penicillium oxalicum*. Biotechnol Biofuels. 2017;10:100.

25. Coradetti ST, Craig JP, Xiong Y, Shock T, Tian C, Glass NL. Conserved and essential transcription factors for cellulase gene expression in ascomycete fungi. Proc Natl Acad Sci. 2012;109:7397–402.

26. Huberman LB, Coradetti ST, Glass NL. Network of nutrient-sensing pathways and a conserved kinase cascade integrate osmolarity and carbon sensing in *Neurospora crassa*. Proc Natl Acad Sci. 2017;114:E8665–74.

27. Raulo R, Kokolski M, Archer DB. The roles of the zinc finger transcription factors XlnR, ClrA and ClrB in the breakdown of lignocellulose by *Aspergillus niger*. AMB Express. 2016;6:1–12.

28. De Souza WR, Maitan-Alfenas GP, de Gouvêa PF, Brown NA, Savoldi M, Battaglia E, et al. The influence of *Aspergillus niger* transcription factors AraR and XlnR in the gene expression during growth in D-xylose, L-arabinose and steam-exploded sugarcane bagasse. Fungal Genet Biol. 2013;60:29–45.

29. Noguchi Y, Tanaka H, Kanamaru K, Kato M. Xylose triggers reversible phosphorylation of XlnR, the fungal transcriptional activator of xylanolytic and cellulolytic genes in *Aspergillus oryzae*. Biosci Biotechnol Biochem. 2014;75:5:953–9.

30. Stricker AR, Grosstessner-Hain K, Würleitner E, Mach RL. Xyr1 (Xylanase regulator 1) regulates both the hydrolytic enzyme System and D-xylose metabolism in Hypocrea jecorina. Eukaryot Cell. 2006;5:2128–37.

31. Miehe H. Die Selbsterhitzung des Heus. Eine biologische Studie. Verlag von Gustav Fischer. 1907.

32. Ruiz-Díez B. A Review: Strategies for the transformation of filamentous fungi. J Appl Microbiol. 2002;92:189–95.

33. Kunitake E, Tani S, Sumitani JI, Kawaguchi T. A novel transcriptional regulator, ClbR, controls the cellobiose- and cellulose-responsive induction of cellulase and xylanase genes regulated by two distinct signaling pathways in *Aspergillus aculeatus*. Appl Microbiol Biotechnol. 2013;97:2017–28.

34. Hynes MJ, Corrick CM, King JA. Isolation of genomic clones containing the amdS gene of *Aspergillus nidulans* and their use in the analysis of structural and regulatory mutations. Mol Cell Biol. 1983;3:1430–9.

35. Coradetti ST, Pinel D, Geiselman GM, Ito M, Mondo SJ, Reilly MC, et al. Functional genomics of lipid metabolism in the oleaginous yeast *Rhodosporidium toruloides*. Elife. 2018; 7: e32110.

36. Kunitake E, Kobayashi T. Conservation and diversity of the regulators of cellulolytic enzyme genes in Ascomycete fungi. Curr Genet. 2017; 63:951–958.

37. Li Z, Yao G, Wu R, Gao L, Kan Q, Liu M, et al. Synergistic and dose-controlled regulation of cellulase gene expression in *Penicillium oxalicum*. PLoS Genet. 2015;11:1–45.

38. Teichert I, Pöggeler S, Nowrousian M. *Sordaria macrospora*: 25 years as a model organism for studying the molecular mechanisms of fruiting body development. Appl Microbiol Biotechnol. Applied Microbiology and Biotechnology; 2020;3691–704.

39. Goosen T, Bloemheuvel G, Gysler C, Bie DA De, Brock HWJ Van Den, Swart K. Transformation of *Aspergillus niger* using the homologous orotidine-5’-phosphate-decarboxylase gene. Curt Genet. 1987;499–503.

40. Lacroute F. Regulation of Pyrimidine Biosynthesis in *Saccharomyces cerevisiae*. J Bacteriol. 1968;95:824–32.

41. Doudna J a., Charpentier E. The new frontier of genome engineering with CRISPR-Cas9. Science. 2014; 346:1258096.

42. Idnurm A, Urquhart AS, Vummadi DR, Chang S, Van de Wouw AP, López-Ruiz FJ. Spontaneous and CRISPR/Cas9-induced mutation of the osmosensor histidine kinase of the canola pathogen *Leptosphaeria maculans*. Fungal Biol Biotechnol. 2017;4:1–12.

43. Kujoth G, Sullivan T, Merkhofer R, Lee T-J, Wang H, Brandhorst T, et al. CRISPR/Cas9-Mediated Gene Disruption Reveals the Importance of Zinc Metabolism for Fitness of the Dimorphic Fungal Pathogen *Blastomyces dermatitidis*. mBio 2018; 9:e00412–18.

44. Nødvig CS, Nielsen JB, Kogle ME, Mortensen UH. A CRISPR-Cas9 system for genetic engineering of filamentous fungi. PLoS One. 2015;10:1–18.

45. Sun J, Glass NL. Identification of the CRE-1 cellulolytic regulon in *Neurospora crassa*. PLoS One. 2011;6.

46. Murakoshi Y, Makita T, Kato M. Comparison and characterization of α -amylase inducers in *Aspergillus nidulans* based on nuclear localization of AmyR. Appl Microbiol Biotechnol. 2012;1629–35.

47. Huberman LB, Liu J, Qin L, Glass NL. Regulation of the lignocellulolytic response in filamentous fungi. Fungal Biol Rev. 2016;30:101–11.

48. Ebbole DJ. Carbon Catabolite Repression of Gene Expression and Conidiation inNeurospora crassa. Fungal Genet Biol. 1998;25:15–21.

49. Leynaud-Kieffer L, Curran SC, Kim I, Magnuson JK, Gladden JM, Baker SE, et al. A new approach to Cas9-based genome editing in *Aspergillus niger* that is precise, efficient and selectable. PLoS One. 2019;1–13.

50. Bistis GN. Chemotropic Interactions Between Trichogynes and Conidia of Opposite Mating-Type in Neurospora Crassa. Mycologia. 1981;73:959–75.

51. Braus GH, Krappmann S, Eckert SE. Sexual development in ascomycetes. Fruit body formation of *Aspergillus nidulans*. Mol Biol Fungal Dev. 2002;215–244.

52. Aguilar-Pontes MV, Zhou M, Van Der Horst S, Theelen B, De Vries RP, Van Den Brink J. Sexual crossing of thermophilic fungus *Myceliophthora heterothallica* improved enzymatic degradation of sugar beet pulp. Biotechnol Biofuels. 2016;9:1–14.

53. Van Den Brink J, Samson RA, Hagen F, Boekhout T, De Vries RP. Phylogeny of the industrial relevant, thermophilic genera *Myceliophthora* and *Corynascus*. Fungal Divers. 2012;52:197–207.

54. Seidl V, Seibel C, Kubicek CP, Schmoll M. Sexual development in the industrial workhorse *Trichoderma reesei*. Proc Natl Acad Sci. 2009;106:13909–14.

55. Dahlmann T, Böhm J, Becker K, Kück U. Sexual recombination as a tool for engineering industrial *Penicillium chrysogenum* strains. Curr Genet. 2015;61.

56. Westergaard M, Mitchell HK. *Neurospora*. V. A synthetic medium favoring sexual reproduction. Am J Bot. 1947;35:573–7.

57. Abràmoff MD, Magalhães PJ, Ram SJ. Image processing with imageJ. Biophotonics Int. 2004;11:36–41.

58. Concordet JP, Haeussler M. CRISPOR: Intuitive guide selection for CRISPR/Cas9 genome editing experiments and screens. Nucleic Acids Res. 2018;46:W242–5.

