## Supplemental Information for "Development of genetic tools for the thermophilic filamentous fungus *Thermoascus aurantiacus*"

Supplemental Tables 1-4

Supplemental Figures 1-2

**Supplemental Tables**

**Supp. Table 1**: List of plasmids used in this study

| Name | Bacterial marker | fungal marker | Parent Plasmid | Insert | JBEI Registry |
|---|---|---|---|---|---|
| pTS57 | Kan | *hph* | NA | *gfp* | JPUB_017131 |
| pTS67 | Kan | *hph* | pTS57 | *xlnR* | JPUB_017129 |
| pJP1 | Kan | *hph* | pTS57, JP36_1 | *cas9*, *gRNA 1* | JPUB_017147 |
| pJP3 | Kan | *hph* | pTS57, JP36_3 | *cas9*, *gRNA 3* | JPUB_017149 |

**Supp. Table 2**: List of strains derived from *T. aurantiacus* ATCC 26904 used in this study

| Name | Genotype | Phenotype | JBEI Registry ID |
|---|---|---|---|
| hph-1 | *hph* ectopic integration | Resistance to hygromycin B | JPUB_017144 |
| hph-2 | *hph* ectopic integration | Resistance to hygromycin B | JPUB_017145 |
| taRG003 | *xlnR* ectopic integration | Xylanase hyperproduction | JPUB_017132 |
| taRG005 | *xlnR* ectopic integration | Xylanase hyperproduction | JPUB_017133 |
| taRG010 | *xlnR* ectopic integration | Xylanase hyperproduction | JPUB_017134 |
| taRG015 | *xlnR* ectopic integration | Xylanase hyperproduction | JPUB_017135 |
| taRG020 | *xlnR* ectopic integration | Xylanase hyperproduction | JPUB_017136 |
| taRG023 | *xlnR* ectopic integration | Xylanase hyperproduction | JPUB_017137 |
| taRG025 | *xlnR* ectopic integration | Xylanase hyperproduction | JPUB_017138 |
| taRG027 | *xlnR* ectopic integration | Xylanase hyperproduction | JPUB_017139 |

| | | | |
|---|---|---|---|
| taRG028 | *xlnR* ectopic integration | Xylanase hyperproduction | JPUB_017140 |
| taRG029 | *xlnR* ectopic integration | Xylanase hyperproduction | JPUB_017141 |
| FOAR1 | UV-induced *pyrE* insertion | 5-FOA resistant, uracil auxotroph | JPUB_017142 |
| FOAR2 | UV-induced *4yre* SNP | 5-FOA resistant, uracil auxotroph | JPUB_017143 |
| FOAR2 *x* taRG008 | Sexual cross of *4yre* SNP and *hph* insertion | 5-FOA resistant, uracil auxotroph, hygromycin B resistant | JPUB_017156 |
| JP1-1 | Cas9-induced pyrG mutation | 5-FOA resistant, uracil auxotroph | JPUB_017150 |
| JP1-2 | Cas9-induced pyrG mutation | 5-FOA resistant, uracil auxotroph | JPUB_017151 |
| JP1-3 | Cas9-induced pyrG mutation | 5-FOA resistant, uracil auxotroph | JPUB_017152 |
| JP1-4 | Cas9-induced pyrG mutation | 5-FOA resistant, uracil auxotroph | JPUB_017153 |
| JP1-5 | Cas9-induced pyrG mutation | 5-FOA resistant, uracil auxotroph | JPUB_017154 |
| JP1-6 | Cas9-induced pyrG mutation | 5-FOA resistant, uracil auxotroph | JPUB_017155 |

**Supp. Table 3**: List of primers used in this study

| PCR | | name | Sequence |
|---|---|---|---|
| *hph1* | FWD | RG1 | CTCGGAGGGCGAAGAATCTC |
| | REV | RG2 | ATTTGTGTACGCCCGACAGT |
| *hph2* | FWD | TS222 | CGTAGTACCTGAGCACCCCTCTGAGCTCTT |
| | REV | TS223 | CCATTTGTCTCAACTCCGGAGCTGACATCGA |
| *pyrE* | FWD | RG75 | GACGGTTTCTATACAGTCTTTTCAG |
| | REV | RG76 | CCCCCGATGTTACTCCGC |
| *pyrG* | FWD | LLK683 | TTCTTACTACAACTTGGCAACCTTC |
| | REV | LLK686 | ACAAGCCAAATTACCAGCAGAATAC |

**Supp. Table 4**. List of protospacers and PAM sequences used in this study

| **Target locus** | **ID** | **Protospacer sequence (5'-3')** | **PAM (5'-3')** |
|---|---|---|---|
| *pyrG* | gRNA 1 | CTTTTGCGCGCGAGCGCCGT | AGG |
| *pyrG* | gRNA 2 | GAGTCTTCCTGCACAGGCCT | GGG |
| *pyrG* | gRNA 3 | TCGGCGCCCGACTTCCCCTA | CGG |

**Supplemental Figures**

**Supp. Figure 1**

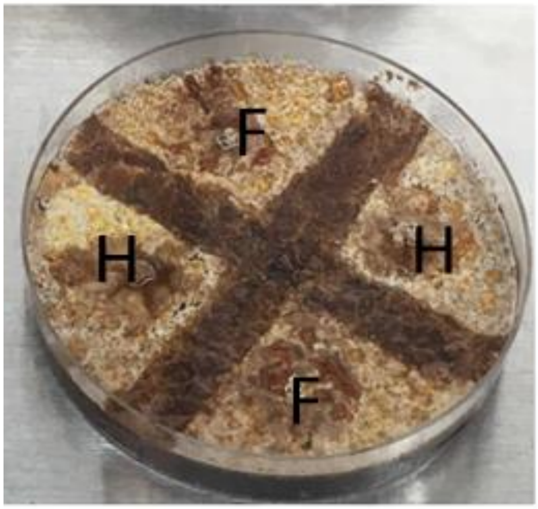


**S. Fig. 1:** Outcrossing of *T. aurantiacus*. Image of the plate setup for strain crossings: the two parent strains were plated in alternating fashion (F: 5-FOA resistant parent strain FOAR2, and H: hygromycin B resistant parent strain taRG008).

**Supp. Figure 2a,b**

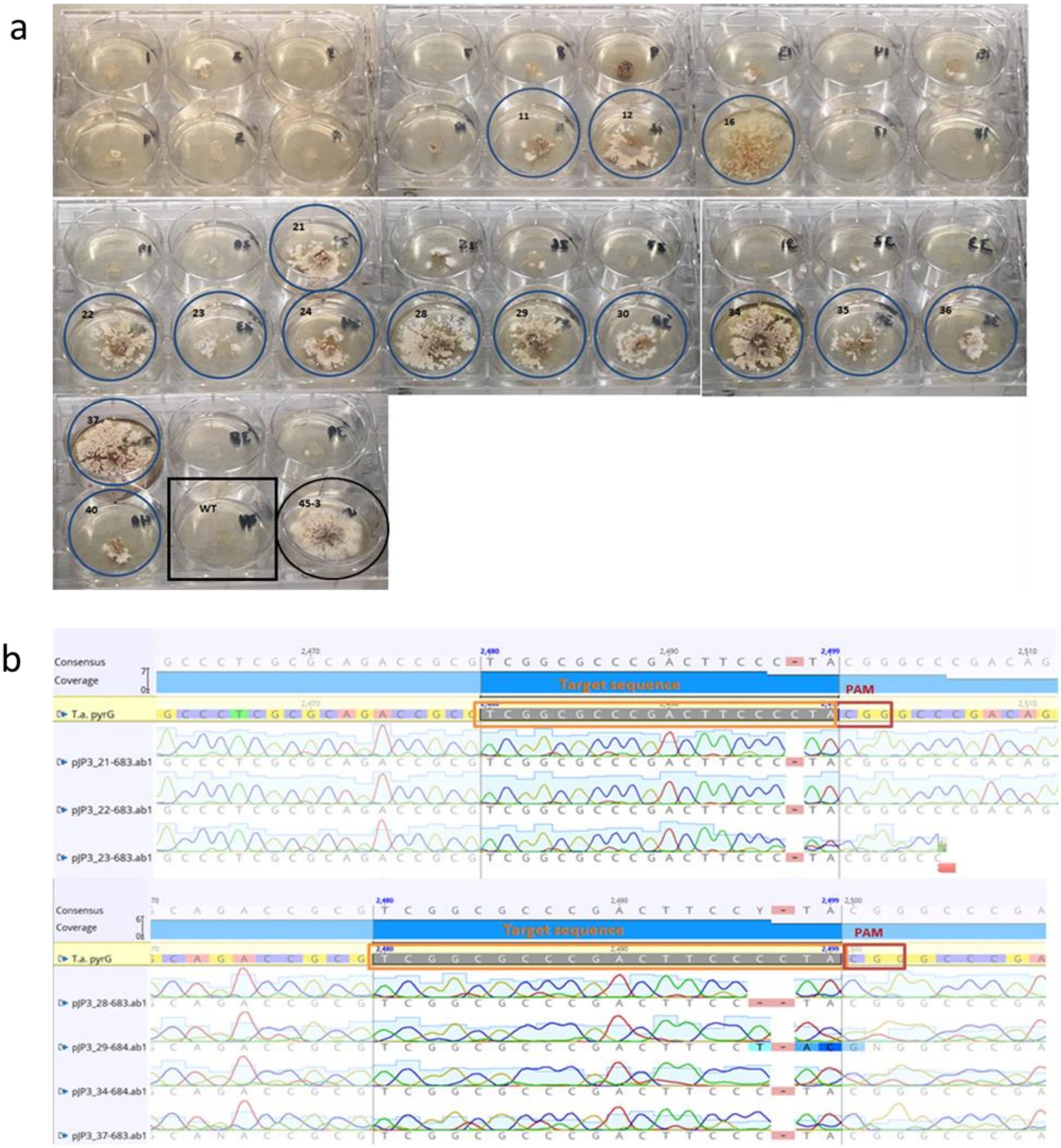

**S. Fig. 2:** Screening for *pyrG* deletion strains on 5-FOA uracil medium. (a) A subset of 20 colonies from ATMT transformations using pJP1 (gRNA 1) and pJP3 (gRNA 3) were selected for 5-FOA resistance each. The wild type (black square frame) and FOAR2 as a 5-FOA resistant positive control (black round frame) were included. pJP1 colony 11, 12 and 16 as well as pJP3 colony 21, 22-24, 28-30, 34-37 and 40 were positive transformants on the selection medium and were used for Sanger sequencing verification procedures. (b) Sanger sequencing results for *T. aurantiacus* pJP3 transformants revealing deletions and mismatches through Cas9 cleavage next to the PAM sequence (framed in red) in the *pyrG* target sequence (framed in orange). The sequence analysis was performed with Geneious version 11.1 (Biomatters).
